# BOOPTHAT: An inexpensive and scalable system for spatiotemporal activation of heat-shock transgenes in zebrafish

**DOI:** 10.1101/2025.06.05.658177

**Authors:** Darren Wang, Benjamin L. Martin

## Abstract

We present the BOOPTHAT (Batch-Operating Optically Powered Targeted Heater for Activating Transgenes) as a low-cost system for activating heat-shock inducible transgenes with spatiotemporal control in multiple zebrafish embryos at a time. Gaining finer spatiotemporal control over gene expression is critical for unraveling the regulatory networks that coordinate embryogenesis. While the heat-shock inducible gene expression system is a widely used tool for controlling temporal transgene expression, its applicability in spatiotemporal control is limited. By adding a level of spatial control onto this well-established system, we take advantage of the existing infrastructure surrounding the HS induction system and introduce new ways to use the multitude of existing lines. The BOOPTHAT system is built from 3D printed components and inexpensive consumer parts. Independent 3D printed micromanipulators are used to position optical fiber probes. When coupled to a light source, the probes are heated photothermally and are used to perform targeted gene activation in multiple samples at a time. We demonstrate the capabilities of our system and highlight some areas of research that stand to benefit from this frugal and effective system.

## Introduction

Genes play different roles throughout the course of development influenced by timing, the location and tissue type that the gene is induced in, and interactions with various genetic pathways. Developing tools to perturb these pathways has been integral to the field of developmental biology. Many commonly used genetic perturbation techniques afford temporal control over global gene expression using inducible gene expression systems, but spatial control is more limited. Tissue level control is achievable with tissue specific promoters like the Gal4-UAS system and tissue-directed transplants. (Halpern et al., 2008)

The *hsp70l* heat shock promoter is a widely used inducible promoter in zebrafish (Bajoghli et al., 2004; Choe et al., 2021). Many HS-inducible fish lines have already been created, making it desirable to directly add a level of spatial control onto this existing infrastructure.

Spatiotemporal control can be achieved through the HS gene induction system by targeting a specific region of the transgenic fish for heating. The difficulty lies in making a reliable and maneuverable heating element at the scale relevant for spatial control in the sub-millimeter scale embryos.

Prior approaches for achieving spatiotemporal control over heat-shock inducible transgenes can be split into two clades 1) the physical heater element approach where a small hot object is placed in contact with the specimen and 2) the laser-based approach where focused light is used to activate the transgene either through direct photothermal heating of tissue or from phototoxicity related stress. Early attempts to use a physical heater to activate heat-shock transgenes included the use of a heated needle in drosophila and a sharpened soldering iron for use in zebrafish (Hardy et al., 2007; Monsma et al., 1988). These physical systems have limitations in spatial control and throughput. Laser-based approaches are promising because of the techniques available for manipulating light at cellular scales but are expensive and can face difficulties with phototoxicity and temperature calibration (Deguchi et al., 2009.; Halloran et al., 2000; Shoji & Sato-Maeda, 2008).

In 2009, Placinta et al. introduced an inexpensive heater system that operated by coupling laser light through an optical fiber to direct the light to the desired location where it was absorbed by a dark coating at the end of the fiber (Placinta et al., 2009). This resulted in an easily maneuverable and localized heat source that could be used to activate heat-shock transgenes in targeted locations. Fibers could be drawn to different diameters using a micropipette puller to achieve different activation area sizes or to activate subcutaneous tissue with thinner fibers. The major limitation to their system is the low throughput. In the described setup, embryos are processed serially with each embryo requiring ∼30 minutes to heat shock. Directly parallelizing the setup to improve throughput would be impractical because a) it used a costly optomechanical system to couple laser light into the optical fiber and b) would require a microscope per replicate setup. The limited throughput would lengthen experiments and make it difficult to apply the system in experiments that require controlled staging of embryos.

We sought to build a system using the same photothermal approach but with increased throughput and convenience. Figure 1 shows the concept of photothermal heating and the general layout of the design that we arrived at.

**Figure 1.**
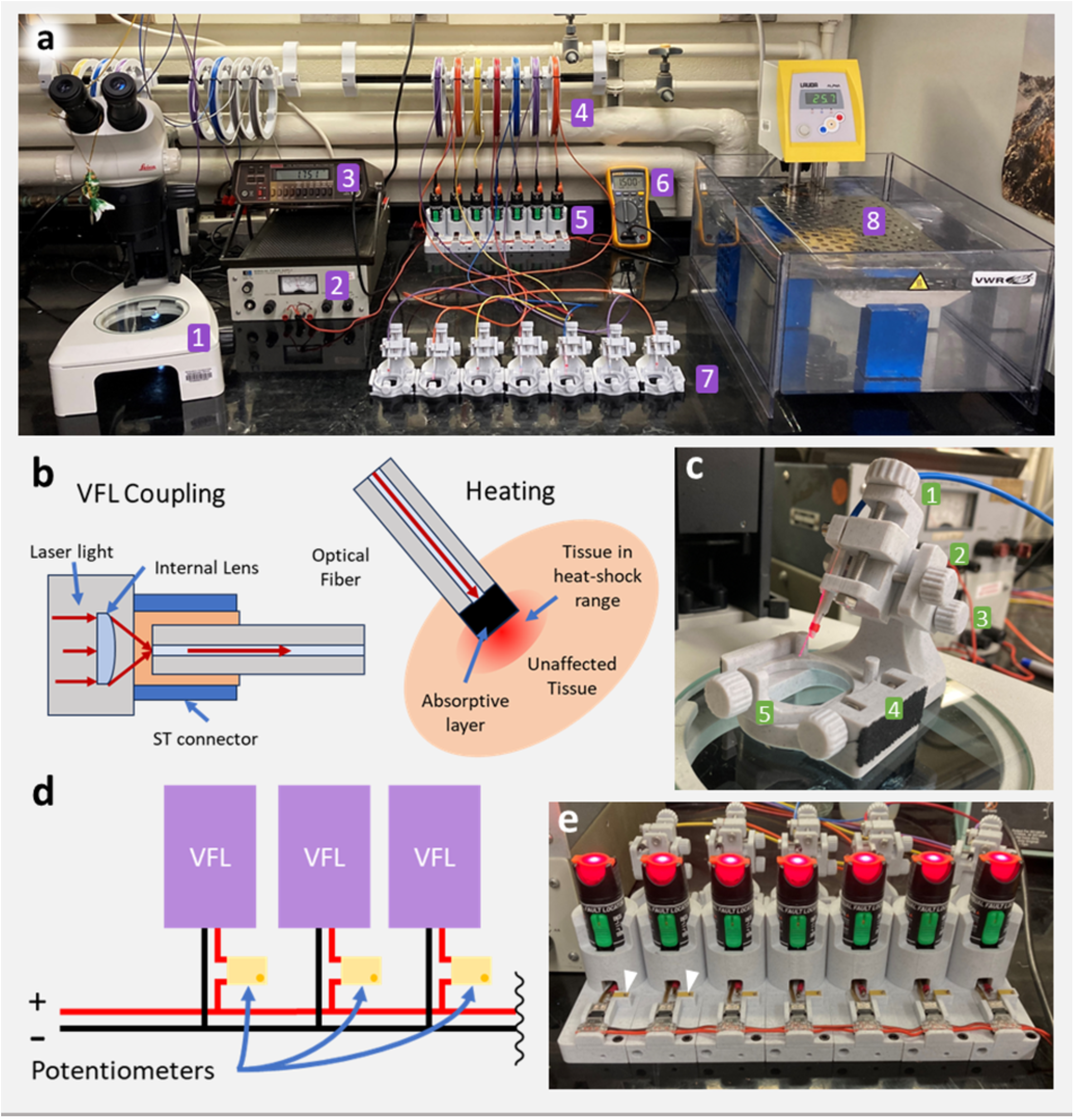
a) full BOOPTHAT setup: 1 - dissection microscope for aligning the micromanipulator probes, 2 - power supply, 3 - voltmeter, 4 - fiber reels on reel rack, 5 - VFL array, 6 - ammeter, 7 - 3d printed micromanipulator cassettes, 8 - temperature controlled water bath for controlling the ambient temperature at which the shock takes place. b) schematic showing the coupling of the VFL with the optical fiber and the photothermal heating probe. c) closeup on one of the 3d printed micromanipulator cassettes, currently turned on, fiber visibly glowing from the small amount of light reflected off the absorptive coating. 1 - extension rail, 2 - y axis rail, 3 - pivot, 4 - x-axis rail for moving petri dish holder, 5 - petri dish holder set screw. d) wiring schematic of the VFL array showing parallel wiring and the positioning of the potentiometers. e) closeup of the VFL array turned on without the optical fibers plugged in, caps on.

**Figure 2.**
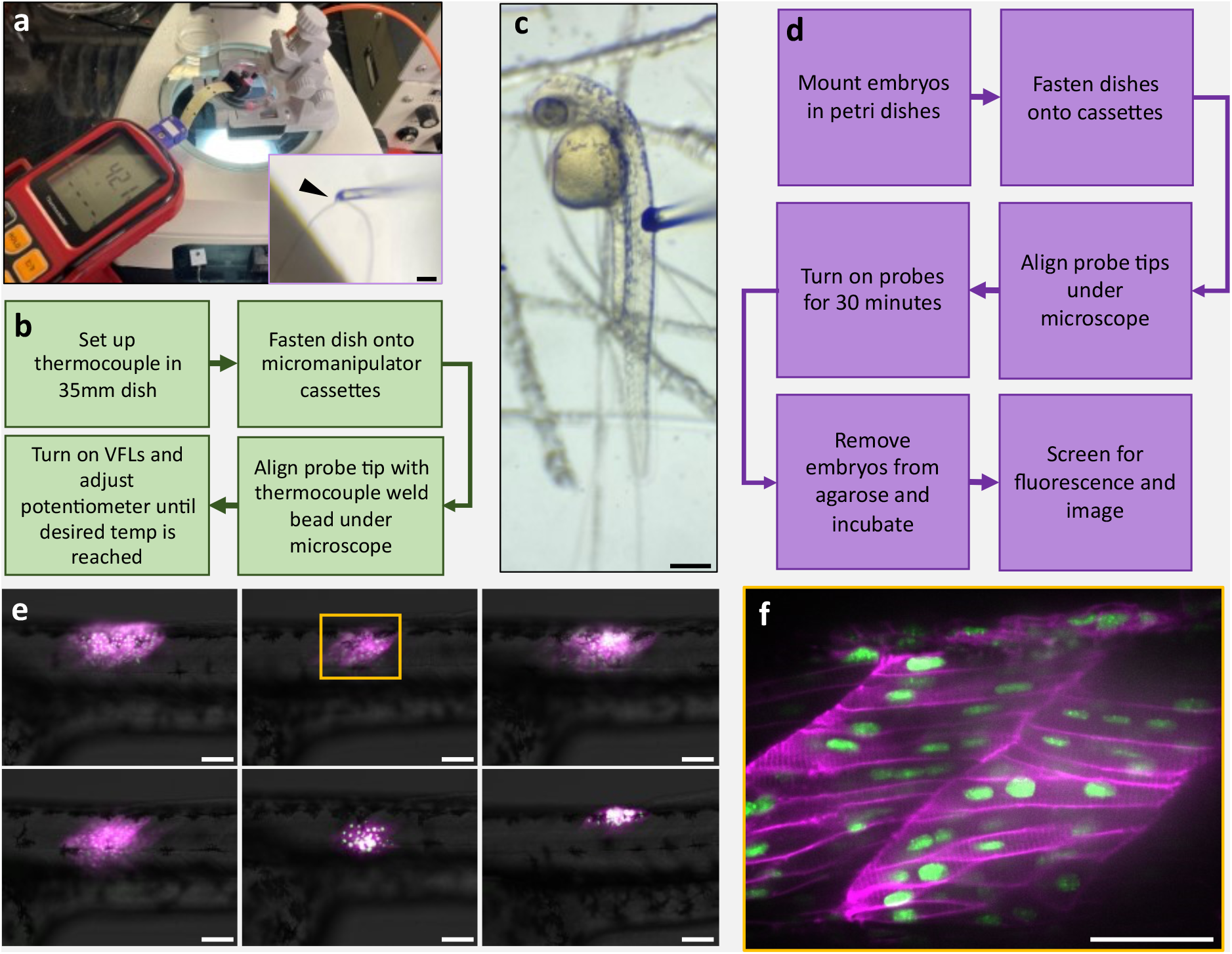
a) thermocouple setup for tuning probe temperature. 15 AWG wire welded to form a bead at the end held in place by a custom 3d printed holder in a petri dish embedded in low melt agarose. Inset: probe tip brought in contact with thermocouple bead, arrow pointing at contact point, 250um scale bar. The temperature of each probe is adjustable using the potentiometer in series with each VFL. b) Workflow for tuning the thermocouples. c) coated end of probe placed in contact with a 36hpf *hsp70l:CAAX-mCherry-P2A-NLS-KikGR* embryo mounted in low melt agarose, scratches in petri dish meant to help agarose droplets adhere, 250um scale bar. d) workflow for booping embryos. e) Images taken from a single batch of embryos booped at ∼40°C probe temperature, 21°C ambient temperature, levels adjusted for clarity. KikGR mapped to green, CAAX-mCherry mapped to magenta, 100 um scale bar. f) confocal image of booped region, 50 um scale bar).

In what follows, we characterize our system and demonstrate how it opens new opportunities investigating developmental processes. Our setup is designed to be easily replicable, with parts being either 3D printable or readily available as consumer goods (Detailed build instructions and bill of materials in GitHub repository linked in supplement). We hope that this setup can bring new ways to use existing reagents to address unanswered questions.

## Results & Discussion

### An inexpensive and scalable optical fiber microheater system

The BOOPTHAT setup consists of multiple independent 3d printed micromanipulator modules, each with an attached petri dish holder. The arrangement allows the modules to be moved around even after positioning the probes on the embryo without losing alignment. Each module can thus be sequentially aligned with a specimen under a dissection microscope and moved to the side before power is turned on for all the modules. To streamline and delineate references to the process of using this system, we humbly offer a new verb for addition to the nomenclature: “to boop.”

Each module is powered by a 30mW, 650nm visual fault locator (VFL), a telecommunications tool used to locate breaks in optical fiber. The VFL can be easily coupled to optical fibers using standard connectors, making it unnecessary to have a separate optomechanical coupling setup. In the BOOPTHAT setup, the VFLs are connected in parallel to a benchtop power supply with a trimmer potentiometer wired in series with each VFL to provide independent power adjustment to each probe. With each VFL connected to an independent micromanipulator cassette, the number of modules is in principle only limited by the capacity of the power supply and desk space.

During construction of the system, we found that different brands of VFLs have different efficiencies in converting electrical power to optical power as well as different current/voltage relationships. During prototyping, the VFL used had an 18% power efficiency but the different (slightly cheaper) VFLs used for the final setup only had a 6% power efficiency with a different current/voltage relationship. Because of this inconsistency, some characterization is needed when replicating this setup. (See supplement for details).

The fiber reel system makes it convenient to switch between different probe in case a damaged probe needs to be replaced, or a different probe diameter is needed for a particular experiment (Fig 1(a)). Fibers can be tapered and cut to achieve thinner diameters using either the flame of a Bunsen burner or the Sutter micropipette puller, as Placinta et al. reported. By having fibers in long excess, rebuilding broken fibers is less wasteful. Most of our testing was done using unpulled fibers but it was found that thinner fibers work as reported by Placinta et.al. (data not shown).

In place of the sharpie pro coating used by Placinta et al., it was found that using black 4.0, an ultra-black acrylic paint, mixed with black acrylic primer in around a 1:1 ratio gave a more durable coating that was able to absorb more of the incident light. With this coating, light leakage from the tip of the fiber was undetectable although there was a minor amount of reflection back up the fiber. We found that the acrylic paint coating did not adhere well to fibers thinner than around 50um and that it was better to use the sharpie pro coating used by Placinta et al. in these situations.

Using a 56 AWG (∼14um) diameter E-type thermocouple embedded in 0.5% low melt agarose to mimic the environment of an agarose-mounted embryo, we were able to measure the temperature of the probe tip. The conduction of heat through the thermocouple wires was approximated as negligible. Consistent with Placinta et al. ‘s findings, we observed that a VFL optical power output in the range of 10mW was enough to achieve temperatures relevant for heat-shock response activation in maximum diameter uncut fibers.

We tested using our system on embryos ranging from 10hpf to 48hpf in age. In preparation for booping, embryos were anesthetized with tricaine and mounted in 0.5% low-melt agarose in small wells made in a thin layer of 1% low-melt agarose in 35mm petri dishes. The process was sped up by setting the agarose in an 18°C incubator. Multiple embryos could be mounted per dish to further increase throughput.

Characterization of the system was done using 36 hpf *Tg(hsp70l:CAAX-mCherry-P2A-NLS-KikGR)* zebrafish embryos (Goto et al., 2017). Before setting up the embryos for booping, each probe was tuned to the desired temperature using the thermocouple embedded in low melt agarose. After embryos were mounted in the petri dishes and the dishes were loaded onto the cassettes, each cassette was adjusted under a dissection microscope to place the probe onto the target region of the embryo one at a time. Once all the microheater cassettes were aligned, the system was turned on for 30 minutes. After shutting off the system and retracting the probes, the embedded embryos were freed from the agarose by cutting a cross shape in the agarose with the tip off a hypodermic needle centered around the embedded embryo. The embryos were left to develop until they reached the desired developmental stage for imaging.

We saw reliable activation (>80% of samples) of heat shock transgenes at all developmental stages that were tested. The general size of the activated areas was consistent when the same conditions were used. We observed some variation, possibly due to probe misalignment when shifting the cassettes around or from uneven heat distribution in the tip due to a poorly applied coating layer resulting in temperature heterogeneity.

The activation tended to have a sharp cutoff laterally at the end of the somite, likely a result of the syncytial nature of the muscle cells and the physical separation of cells between adjacent somites.

### Altering ambient temperature affects the size and shape of activated region

With the idea of gaining better control over properties of the booped area, we added a temperature-controlled water bath used to modulate the ambient temperature (figure 1). The cassettes were placed so that the petri dishes were surrounded by the temperature-controlled water. By adjusting the difference between the heat-shock temperature and the ambient temperature it was possible to alter the temperature gradient and change the geometry of the booped area. All probes were tuned at room temperature for convenience but those being placed in the water bath were tuned to a lower temperature to compensate for the elevated temperature of the water bath. We calculated this temperature by subtracting the difference between the water bath temperature and the ambient temperature in which the probes were tuned from the target probe temperature. In our tests, the water bath temperature was 28°C, the room temperature was 21°C, and the target probe temperature was 40°C so we tuned our probes to 40-(28-21) = 33°C outside of the water bath.

36 hpf *Tg(hsp70l:CAAX-mCherry-P2A-NLS-KikGR)* were booped and imaged around the 48 hpf stage. Embryos booped at 28°C ambient temperature had noticeably larger regions of activation in comparison to those booped at 21°C (fig 3, b).

**Figure 3.**
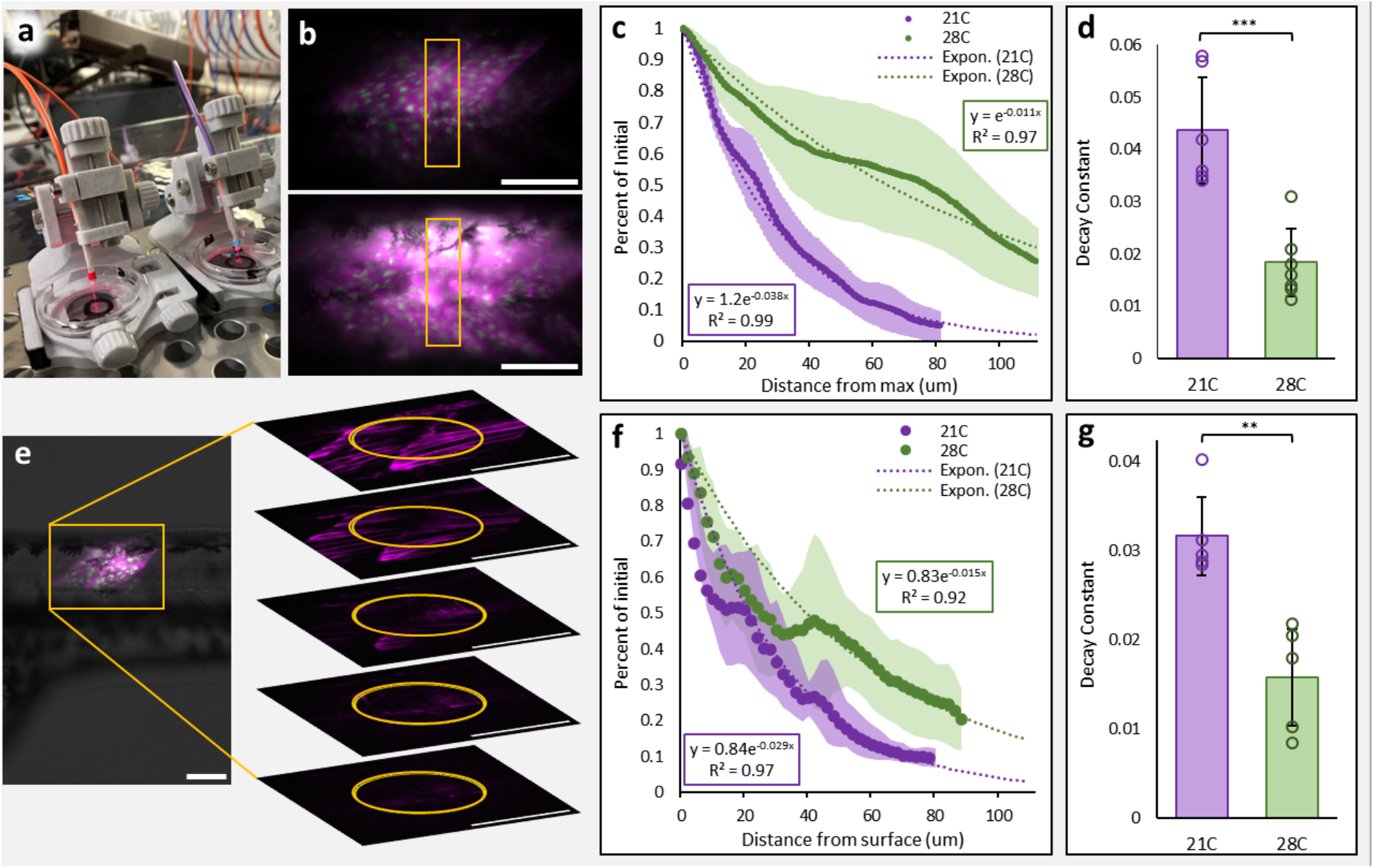
a) closeup view of the micromanipulator cassette placed in the water bath, water is in contact with the sides of the petri dish. b) view of activated region in an embryo booped at an ambient temperature of 21°C (top) and 28°C (bottom); the region analyzed for fluorescence intensity decay along the dorsal-ventral axis is boxed in; 100um scale bar. c) graph showing decay of CAAX-mCherry fluorescence intensity along the dorsal-ventral axis for embryos booped at an ambient temperature of 21°C vs 28°C. d) graph comparing the fitted CAAX-mCherry fluorescence intensity decay rate constants along the dorsal-ventral axis of embryos booped at an ambient temperature of 21°C vs 28°C. e) view of a booped region with a closeup showing the fluorescence intensity at various depths along the mediolateral axis with the top picture being the most lateral; the region analyzed for fluorescence intensity decay along the mediolateral axis is circled; 100um scale bar. f) graph showing decay of CAAX-mCherry fluorescence intensity along the mediolateral axis for embryos booped an ambient temperature of 21°C vs 28°C. g) graph comparing the fitted CAAX-mCherry fluorescence intensity decay rate constants along the mediolateral axis of embryos booped at an ambient temperature of 21°C vs 28°C36 hpf *Tg(hsp70l:CAAX-mCherry-P2A-NLS-KikGR)* were booped and imaged around the 48 hpf stage. Embryos booped at 28°C ambient temperature had noticeably larger regions of activation in comparison to those booped at 21°C (fig 3, b).

**Figure 4.**
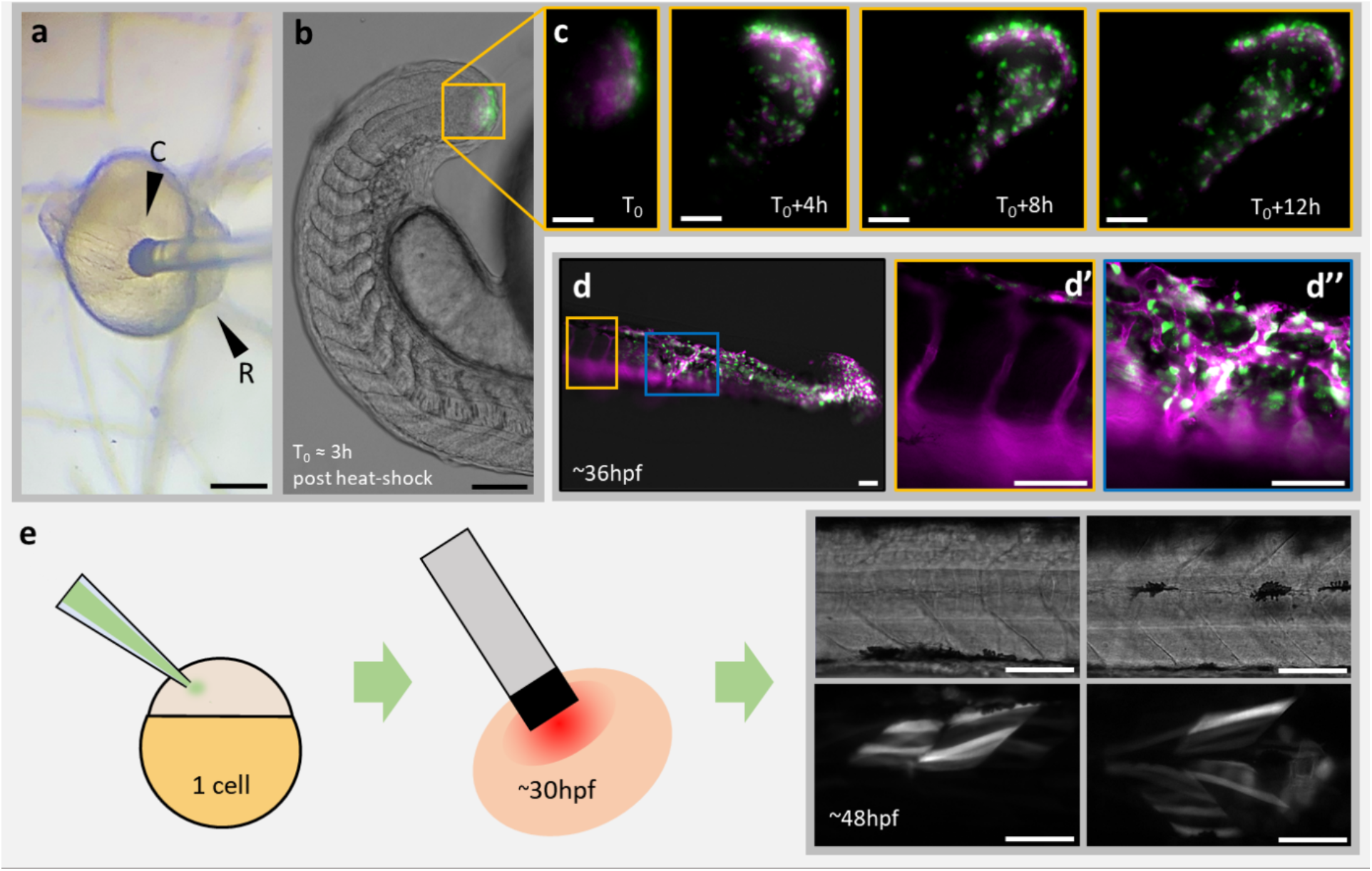
a) 15 hpf *hsp70l:id3-2A-NLS-KikGR* / *kdrl:Hsa*.*HRAS-mCherry* embryo mounted in low melt agarose gel with probe positioned on the tailbud (250um scale bar). b) tailbud booped *hsp70l:id3-2A-NLS-KikGR* / *kdrl:Hsa*.*HRAS-mCherry* embryo ∼3hpb (hours post boop) showing kikGR mapped to green and mCherry mapped to magenta (150um scale bar). c) timelapse of embryo in b showing cells dispersing over time (50um scale bar). d) separate experiment of 17hpf *hsp70l:id3-2A-NLS-KikGR* / *kdrl:Hsa*.*HRAS-mCherry* embryo booped in the presomitic mesoderm showing booped cells contributing vascular endothelium overgrowth (d’’) as opposed to normal endothelium (d’) (50um scale bars). e) Boop of plasmid-injected embryos leads to spatiotemporal mosaic activation of injected plasmid (100um scale bars).

We quantified the fluorescence intensity profile along both the dorsal-ventral axis (fig 3, b-d) and the mediolateral axis (fig 3, e-g). Along the dorsal-ventral axis, we found the average fluorescence intensity starting from the point of max intensity and moving towards the side with the slower rate of decay. We plotted the fluorescent intensity, normalized to the max intensity, and empirically fitted the plot with a decaying exponential function. The decay rate of fluorescent intensity along the dorsal-ventral axis of average intensity projected image stacks was less in embryos booped at 28°C than in those booped at 21°C. (fig 3, c-d). Along the mediolateral axis, we found the average fluorescence intensity for a region surrounding the probe contact point starting from the beginning (lateral) of the muscle and moving medially (fig 3, e). Like before, we plotted the fluorescent intensity, normalized to the max intensity, then empirically fitted the graph with a decaying exponential. As with the dorsal-ventral axis, the decay rate of fluorescent intensity along the medial axis in embryos booped at 28°C was less than for embryos booped at 21°C (fig 3, f-g).

This ability to alter the gradient of transgene activation brings up the possibility of manipulating morphogen gradients using heat-shock inducible copies of cell-autonomous downstream factors or inhibitors in morphogen signaling pathways.

### Mosaic transgene expression

Mosaic genetic perturbation is useful for isolating the effects of the perturbation by observing the behavior of modified cells placed in a wild-type environment. One technique to achieve this in zebrafish is by performing tissue-directed transplants using cells from a donor embryo that is genetically distinct from the host embryo (Kemp et al. 2009). Referencing the fate map of the zebrafish embryo, cells from a donor can be transplanted into a region fated to become the desired tissue (Kimmel et al. 1990). Using donors with a HS-inducible version of the gene of interest (GOI), the timing of transgene expression can be controlled by shocking the host embryo at the time point of interest.

As an embryo develops and the extracellular matrix matures, it becomes progressively more difficult to successfully transplant cells. This limits transplants to the initial stages of embryonic development which in turn limits control over the resulting distribution of transplanted cells.

Transplants are also a difficult procedure that requires precise scheduling, fine motor skills, and experience to pull off correctly.

We found two ways to achieve mosaic gene expression with the BOOPTHAT. One method resembles transplants – a patch of tissue was booped at an earlier stage and the cells spread out over time to create a mosaic distribution. Compared to transplants, the initial induction could be performed later in development, giving more control over the final distribution of cells. This method also had a more streamlined protocol than transplants as it only required embryos from a single HS-inducible fish line. A downside to this approach is not being able to directly control when the cells begin to express the GOI after the boop as can be done with tissue directed transplants using a HS-inducible donor line.

To demonstrate the ability of the system to genetically perturb a specific population of cells, we booped the tailbud and presomitic mesoderm of embryos with both the *hsp70l:id3-2A-NLS-KikGR* and *kdrl:Hsa*.*HRAS-mCherry* transgenes around 15 hpf (Row et al., 2018; Chi et al., 2008). The cells spread out over time and eventually contributed to an overgrowth of vasculature seen around 36hpf, as expected from previous experiments activating *id3* expression (Row et al., 2018).

Another way we were able to achieve mosaic expression was by taking advantage of the mosaic uptake of injected plasmids and booping embryos injected with plasmids containing a HS-inducible version of the GOI. The embryos were injected at the 1 cell stage and left to grow up to the desired stage. When booped, only cells that had taken up the plasmid within the booped region were activated, resulting in spatially controlled mosaic activation of the GOI.

## Conclusion and Outlook

We have presented an inexpensive and scalable microheater setup for activating transgenes under regulation of the *hsp70l* promoter with control over the size and gradient of activation. With this approach, one can achieve reliable and high-throughput spatiotemporal control over any HS-inducible GOI in developing zebrafish embryos. We demonstrated the performance of the system in several different ways. First, we showed that a local population of cells can be labeled with a reporter and that the dimension of transgene activation can be modulated by adjusting the ambient temperature of the embryos when heat-shocked. Second, we demonstrated that the system can be used for mosaic labeling of cells. And third, we showed that we can locally modify the development of tissues by inducing a small patch of ectopic vasculature in a region normally fated to become skeletal muscle.

The ability to activate gene expression in localized groups of cells brings up the exciting possibility of manipulating the morphogen gradients that coordinate development, either by directly expressing a secreted morphogen or morphogen inhibitor in a patch of tissue, or by using the temperature gradient to generate a proxy gradient of cell-autonomous downstream targets of morphogen activity. This use of the BOOPTHAT system could help tease apart the molecular mechanisms governing differential cellular behavior and fate in response to varying levels of morphogen activity.

## Methods

### Probe Cassette Build

The micromanipulators were designed in Fusion 360. They were sliced with 30% gyroid patterned infill, 0.2mm layer height, 8 layers on the top and bottom, and 4 perimeters. They were printed on with PETG filament using a Prusa mk4 3D printer with a 0.4 mm nozzle. These settings gave rigid parts. Instructions and files for assembly are provided in a GitHub repository (https://github.com/Wang-in-100million/BOOPTHAT).

The rail holding holes were reamed with a 4mm diameter drill bit and fitted with 40mm M4 screws and nuts that worked as lead screws. The nuts were press-fitted into place using a small tabletop vice and a small mallet. Grease was used where needed to make the system operate smoothly.

The visual fault locator array was designed in Fusion 360 and printed in PETG. The handles were removed from the 650nm, 30mW VFLs (FYBOPTWU) and 20 AWG wires were soldered onto the leads so that the VFLs could be powered using a benchtop power supply in place of batteries. The VFLs were placed in parallel to each other. A 10 ohm 25 turn trimmer potentiometer (Bourns) was soldered in series with each VFL to allow for independent power adjustment.

56 gauge E type thermocouple wires (OMEGA) were welded using a pencil lead electrode and a benchtop power supply. The fine wires were held in place on a piece of tape folded over itself using smaller pieces of tape and glue (instructions provided in the GitHub repository).

We used a 125um diameter multimode optical fiber with a 50um core that terminated in a standard ST connector on both sides. The fiber was cut in half to make two probes per fiber. The excess fiber was wound around 3D printed reels. At the end of the fiber the protective jacket was removed with a pair of wire strippers, the protective aramid fiber was cut with a knife, and the second protective jacket was removed with wire strippers. The end of the fiber was scored using a micropipette-cutting ceramic tile (Sutter) and broken off to leave a clean surface perpendicular to the path of the light. The coating was made with a roughly 1:1 ratio of Black 4.0 acrylic paint (Culture Hustle) and black acrylic primer (Vallejo). The coating was applied under a dissection microscope using something thin like a piece of pencil lead. The quality of the coating was tested by turning on the VFLs and looking for light leakage from the tip of the fiber (not to be done under microscope, light leakage can be safely seen by pointing the end of the fiber on a nearby surface).

Links to the components needed to replicate the setup can be found in the bill of materials in the Github repository.

### Zebrafish handling

Zebrafish methods were approved by the Stony Brook University Institutional Animal Use and Care Committee. Transgenic lines used in this study include *Tg(hsp70l:CAAX-mCherry-P2A-NLS-KikGR)*^*sbu104*^, *Tg(hsp70l:id3-2A-NLS-KikGR)*^*sbu105*^, and *Tg*(*kdrl:Hsa*.*HRAS-mCherry)*^*s896*^ (Goto et al., 2017; Row et al., 2018; Chi et al., 2008). Embryos were raised in embryo media at 28°C after initial collection. In preparation for booping, embryos were anesthetized with tricaine, mounted in 0.5% low-melt agarose in small wells made by poking holes in a thin layer of 1% low-melt agarose in 35mm petri dishes. After being booped, embryos were freed from the agarose by cutting a cross shape in the agarose with the tip off a hypodermic needle centered around the embedded embryo. Embryos were imaged using either a Leica DMI6000B inverted microscope or a custom-built spinning disk confocal microscope with a Zeiss Imager A.2 frame, a Borealis modified Yokogawa CSU-10 spinning disc, ASI 150uM piezo stage controlled by an MS2000, an ASI filter wheel, and a Hamamatsu ImageEM x2 EMCCD camera (Hamamatsu C9100-23B). The microscope is controlled with Metamorph microscope control software (V7.10.2.240 Molecular Devices), with laser illumination via a Vortran laser merge controlled by a custom Measurement Computing Microcontroller integrated by Nobska Imaging. Laser power levels were set in Vortran’s Stradus VersaLase 8 software.

## Acknowledgements

We thank members of our lab for helpful discussions.

